# Nanobody-mediated Complement Activation to Kill HIV-infected Cells

**DOI:** 10.1101/2022.06.09.495439

**Authors:** Maria Lange Pedersen, Dennis Vestergaard Pedersen, Mikael Becher Lykkegaard Winkler, Heidi Gytz Olesen, Ole Schmeltz Søgaard, Lars Østergaard, Nick Stub Laursen, Anna Halling Folkmar Andersen, Martin Tolstrup

## Abstract

The complement system which is part of the innate immune response against invading pathogens, represents a powerful mechanism for killing of infected cells. Utilizing direct complement recruitment for complement-mediated elimination of HIV-1-infected cells is underexplored. We developed a novel therapeutic modality to direct complement activity to the surface of HIV-1-infected cells. This bispecific complement engager (BiCE) is comprised of a nanobody recruiting the complement-initiating protein C1q, and single-chain variable fragments of broadly neutralizing antibodies (bNAbs) targeting the HIV-1 envelope (Env) protein. Here, we show that two anti-HIV BiCEs targeting the V3 loop and the CD4 binding site, respectively, increase C3 deposition and mediate complement-dependent cytotoxicity (CDC) of HIV-1 Env expressing Raji cells. Furthermore, anti-HIV BiCEs trigger complement activation on primary CD4 T cells infected with laboratory-adapted HIV-1 strain and facilitates elimination of HIV-1-infected cells over time. In summary, we present a novel approach to direct complement deposition to the surface of HIV-1-infected cells leading to complement-mediated killing of these cells.

## Introduction

HIV-1 continues to be a global pandemic claiming hundreds of thousands of lives every year (World Health Organization, 2021, November 30). Despite the advent of combined antiretroviral therapy (cART), there is still no cure for HIV. Currently, there are several broadly neutralizing antibodies (bNAbs) with different binding motives under investigation in clinical trials (Caskey *et al*, 2019). In vitro and in vivo data show that the antiviral effects of monoclonal antibodies against HIV-1 are augmented by complement (Gauduin *et al*, 1997; Posner *et al*, 1992).

The complement system is central to innate immunity and it plays a crucial role in clearance of infectious microbes, including retroviruses such as HIV-1 (Merle *et al*, 2015; Yu *et al*, 2010). The classical complement pathway is initiated by binding of the C1 complex to the Fc moieties of antigen-bound IgG and IgM. The C1 complex is composed of the pattern recognition molecule C1q and two proteases C1r and C1s. C1q is a bouquet-like structure assembled from six collagenous stems, each with a C-terminal globular head that mediate antibody binding (Thielens *et al*, 2017). The affinity between C1q and a single Fc domain is low, consequently activation of C1q is avidity-driven and requires multivalent binding of C1q to immune complexes (Diebolder *et al*, 2014; Feinstein *et al*, 1986). Potent complement activation is thus dependent on antigen size, density, and geometry (Melis *et al*, 2015). Activation of the C1 complex triggers a proteolytic cascade leading to the formation of highly active enzyme complexes, also known as convertases. One central step in the cascade is the cleavage of complement component 3 (C3) into C3a and C3b. Covalently attached C3b on the activator surface promotes different events: i) C3b on target surfaces further opsonization and phagocytosis by effector cells expressing complement receptors for C3b; ii) At high C3b densities exceeding a certain threshold, C3b deposition results in the assembly of C5b-9, also known as the membrane attack complex (MAC), leading to complement-dependent cytotoxicity (CDC) (Bajic *et al*, 2015).

The HIV-1 envelope (Env) is a homotrimer made up by the glycoproteins gp120 and gp41. Antibodies recognizing the HIV-1 glycoproteins can inhibit viral entry into host cells, thus protecting against HIV infection (Zolla-Pazner, 2004). The classical pathway of complement can be activated by direct binding of C1q to the viral Env (Ebenbichler *et al*, 1991; Thielens *et al*, 2002). Furthermore, the classical pathway is enhanced through C1q binding to immune complexes on the virions (Spear *et al*, 1993; Yu *et al*., 2010). Coating HIV-1 virions with complement components is an important mechanism of neutralization which functionally inhibits binding of infectious virus to target cells (Sullivan *et al*, 1998). Conversely, complement protein mediated opsonization can play a critical role in spread and maintenance of virus as deposited complement components on virions facilitate HIV-1 interaction with complement receptor-bearing target cells such as macrophages, monocytes, dendritic cells and B cells (Huber *et al*, 2011). Several reports demonstrated that the HIV-1 particle is susceptible to CDC (Spear *et al*, 1990; Sullivan *et al*, 1996). Furthermore, in vitro studies showed that plasma from HIV-1 infected individuals can mediate CDC of virions (Huber *et al*, 2006; Aasa-Chapman *et al*, 2005). These data highlight the potential of specific activation of complement against HIV-1 infected cells as a part of an HIV cure strategy.

HIV-1 bNAbs bind the HIV-1 Env and neutralize a wide range of HIV strains (Klein *et al*, 2013). BNAbs target different highly conserved epitopes on the viral Env, including the CD4 binding site, the membrane proximal external region (MPER) of gp41, the gp120/gp41 interface, and the N-glycans of the V1/V2 and V3 loop of gp120 (Mouquet, 2014). Administration of bNAbs can suppress plasma viral load in HIV-1-infected individuals (Caskey *et al*, 2015; Caskey *et al*, 2017). Interestingly, an in vitro study showed that the bNAbs 10-1074 and 3BNC117, binding the V3 loop and the CD4 binding site, respectively, were potent complement activators on HIV-1-infected cells (Dufloo *et al*, 2020).

To date, complement-engaging therapeutics are underexplored, and only few therapeutics directly recruit complement to target surfaces. Recently, nanobodies (Nbs) were developed to modulate the complement system (Muyldermans, 2013; Zarantonello *et al*, 2021). Nbs are derived from the VHH domain of heavy chain-only antibodies found in camelids and composed of a single immunoglobulin domain (Muyldermans, 2013). C1qNb75 is the first described Nb modulator of C1q and was generated by immunization of a llama with C1q. In vitro studies showed that C1qNb75 binds C1q with sub-nanomolar affinity and inhibits activation of the classical pathway (Laursen *et al*, 2020). Thus, Nbs are proposed as potential novel complement modulators that could offer new therapeutic strategies (Muyldermans, 2013; Zarantonello *et al*., 2021) Another study investigated a bispecific antibody, with a Fab binding C1q and another Fab binding to a cell surface target. They showed that the bispecific antibodies were capable of recruiting and activating complement on bacterial surface proteins and B and T cell antigens (Cruz *et al*, 2019). These data together, stress the potential of modulation and redirection of complement to target microbial and cell surfaces.

Here, we describe a novel therapeutic modality to direct complement activity to the surface of HIV-1 Env expressing cells with the aim of eliminating these cells. This bispecific complement engager (BiCE) is composed of a nanobody recognizing C1q, and a single-chain variable fragment of a bNAb binding the HIV-1 Env protein. We hypothesized that this BiCE could efficiently recruit C1q, increase cellular complement deposition and mediate CDC of HIV-1-infected cells.

## Results

### Generation and characterization of two anti-HIV BiCEs

We developed a novel treatment modality to recruit C1q directly to the surface of HIV-infected cells with the aim of triggering the complement cascade and subsequent complement-mediated cell lysis, as an immunotherapy concept (Figure 1A). We designed two anti-HIV BiCEs comprised of one part; C1qNb75, a recently discovered nanobody specific for C1q (Laursen *et al*., 2020), and another part; scFv10-1074 or scFv3BNC117, a single-chain variable fragment of the bNAb 10-1074 or 3BNC117 specific for the HIV-1 Env protein (Mouquet *et al*, 2012; Scheid *et al*, 2011). The two domains were fused using a (GGGGS)_3_ linker, and the molecules were termed scFv10-1074-Nb75 and scFv3BNC117-Nb75, respectively. Theoretical molecular weights of the two BiCEs were 43 kDa for scFv10-1074-Nb75 and 45 kDa for scFv3BNC117-Nb75. The anti-HIV BiCEs were expressed in HEK293F cells and purified using affinity chromatography. Protein purification was confirmed by SDS-PAGE analysis (Figure 1B). Characterization of the BiCEs and verification of complex formation to the globular heads of C1q (C1qGH) was analyzed using size exclusion chromatography (SEC). ScFv10-1074-Nb75 and scFv3BNC117-Nb75 eluted almost exclusively in a single peak at 12 mL and 11.8 mL, respectively (Figure 1C). This was in line with their theoretical molecular weights. A clear shift in elution volume was observed for the mixed scFv10-1074-Nb75+C1qGH and for the mixed scFv3BNC117-Nb75+C1qGH, demonstrating complex formation (Figure 1C). In summary, the data obtained confirmed the creation of two novel anti-HIV BiCEs each with a functional nanobody domain able to bind human C1q.

**Figure 1.**
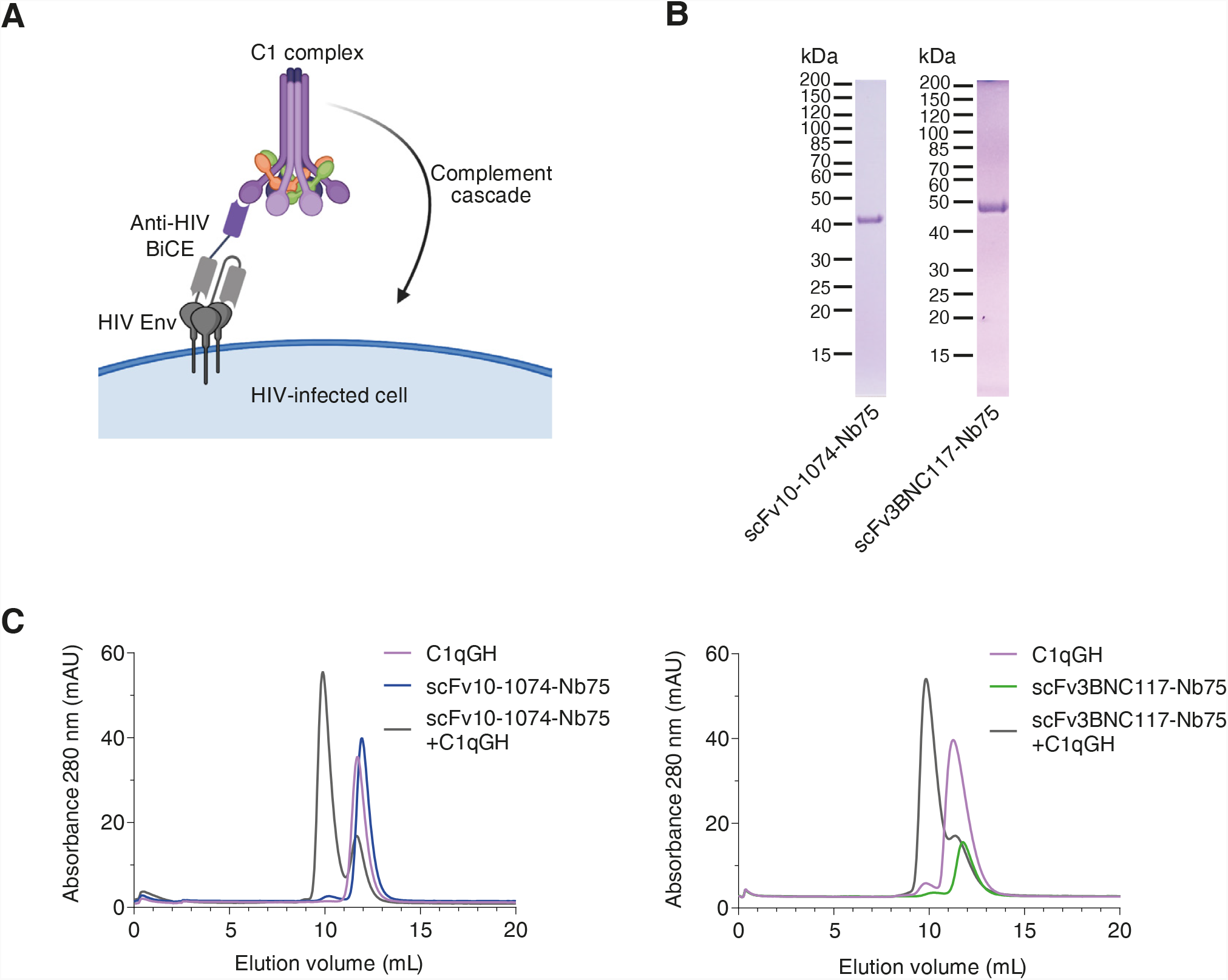
Characteristics of Bispecific Complement Engagers (BiCEs) targeting HIV Env and C1q. A Proposed mechanism of action of the anti-HIV BiCEs. One part of the molecule, a scFv (grey), recognizes the HIV Env. The other part, a nanobody (purple), binds the C1q protein with sub-nanomolar affinity. Engagement of C1q triggers the complement cascade. B Reduced SDS-PAGE analysis of purified anti-HIV BiCEs. C Complex formation using scFv10-1074-Nb75 (43 kDa), scFv3BNC117-Nb75 (45 kDa) and C1qGH (48 kDa) analyzed by SEC on a 24 mL Superdex75 column.

### Characterization of the scFv domain of anti-HIV BiCEs

To determine whether the two novel anti-HIV BiCEs possessed the same HIV neutralizing potency as the full-length monoclonal bNAbs 10-1074 and 3BNC117, we performed a virus neutralization assay. Here, we measured the HIV-infection as HIV Tat-regulated firefly luciferase gene expression in TZM-bl cells after 48 hours of exposure to HIV-1_YU2_. The antibodies or BiCEs were pre-incubated with virus before cells were added to the wells. We observed a concentration-dependent neutralization of HIV from all antibodies or BiCEs, measured by attenuation of luminescence signal with increasing concentrations of antibody or BiCE (Figure 2A). There was a 2-fold difference at the 50% inhibitory concentration (IC_50_) between scFv10-1074-Nb75 (IC_50_ = 0.15 μg/mL) and full-length 10-1074 antibody (IC_50_ = 0.07 μg/mL). Similarly, there was a 9-fold difference in the IC_50_ values between scFv3BNC117-Nb75 (IC_50_ = 0.18 μg/mL) and 3BNC117 (IC_50_ = 0.02 μg/mL), again with the full-length antibody being more potent.

**Figure 2.**
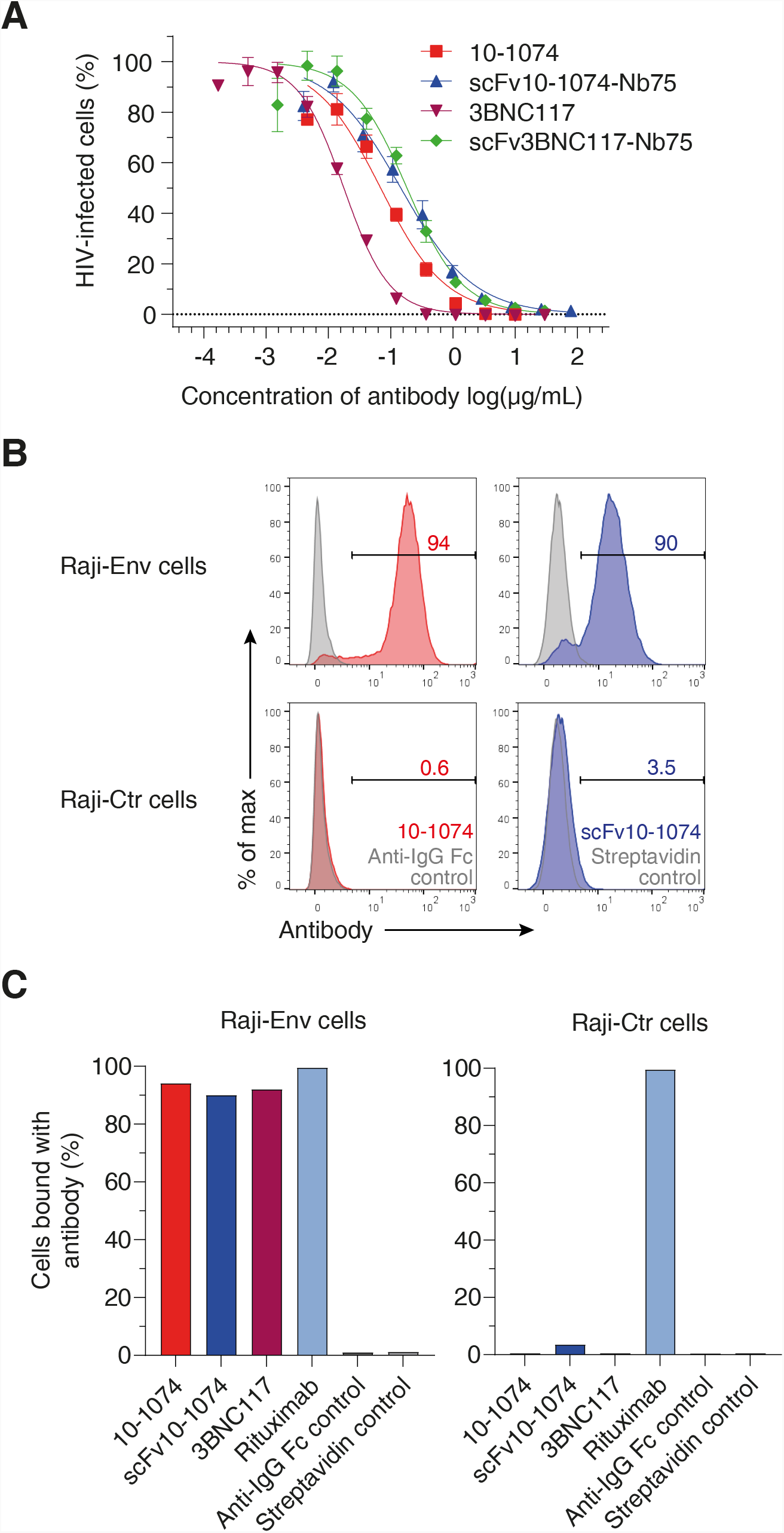
ScFvs domain of anti-HIV BiCEs are capable of recognizing HIV Env and neutralize free virus. A HIV-1_YU-2_ was preincubated with the indicated concentrations of antibodies or BiCEs and subsequently incubated with HIV reporter TZM-bl cells. 48 h later, HIV-infection was measured as luciferase activity quantified by luminescence. For each sample, a normalized HIV-infection is calculated by subtracting RLU of control (no virus added), and then normalized to virus control (no antibody or BiCE added) set to 100%. n = 3 independent experiments, each an average of three independent replicates are shown. B Raji cells expressing HIV-1_YU-2_ envelope (Raji-Env) or Raji control cells (Raji-Ctr) were incubated with 10-1074 or scFv10-1074 or with controls (only secondary antibody or streptavidin added). Binding of 10-1074 and scFv10-1074 were assessed by flow cytometry using a BV421-conjugated anti-IgG Fc antibody or streptavidin, respectively. The numbers indicate the % of cells with bound antibody. One experiment is shown. C Binding of the indicated antibodies to the surface of Raji cells. n = 1 experiment. All data are presented as mean ± SEM.

Next, we used a Raji B cell derivative expressing HIV-1 YU2-Env (Raji-Env cells) to further evaluate Env-recognition and complement activation by our anti-HIV BiCEs. Raji cells express CD20 on the surface, enabling complement activation by rituximab as a positive control (van Meerten *et al*, 2006). To examine the ability and specificity of our antibodies to recognize HIV Env on cell surfaces, we incubated full-length bNAbs and scFv10-1074 with Raji-Env or wildtype Raji control (Raji-Ctr) cells. Antibody recognition was assessed by flow cytometry. More than 90 % of Raji-Env cells were positive for Env (Figure 2B-C). Furthermore, anti-HIV antibodies were not detected on Raji-Ctr cells (Figure 2B-C). Overall, these results show that anti-HIV BiCEs can recognize and neutralize HIV-1 in a concentration-dependent manner, preventing infection of cells. Moreover, the scFv10-1074’s binding ability to Env on cell surfaces was verified.

### Anti-HIV BiCEs trigger complement activation and subsequent lysis of Env-expressing cells

To further explore the anti-HIV BiCEs, we performed a CDC dose-response analysis. We used full-length 10-1074 and 3BNC117 as direct comparison and normal human serum (NHS) as a source of complement. CDC was assessed by measuring cell viability after 24 h incubation at 37°C. Interestingly, the anti-HIV BiCEs displayed a bell-shaped dose-response with highest CDC activity measured at 10 μg/mL (Figure 3A). In contrast, full-length bNAbs plateaued at 10 μg/mL. The reduced effectivity at high BiCE concentration is most likely due to free BiCE molecules competing with BiCE in complex with C1q for binding to Env, thereby preventing recruitment of C1q the cell surface and subsequent CDC.

**Figure 3.**
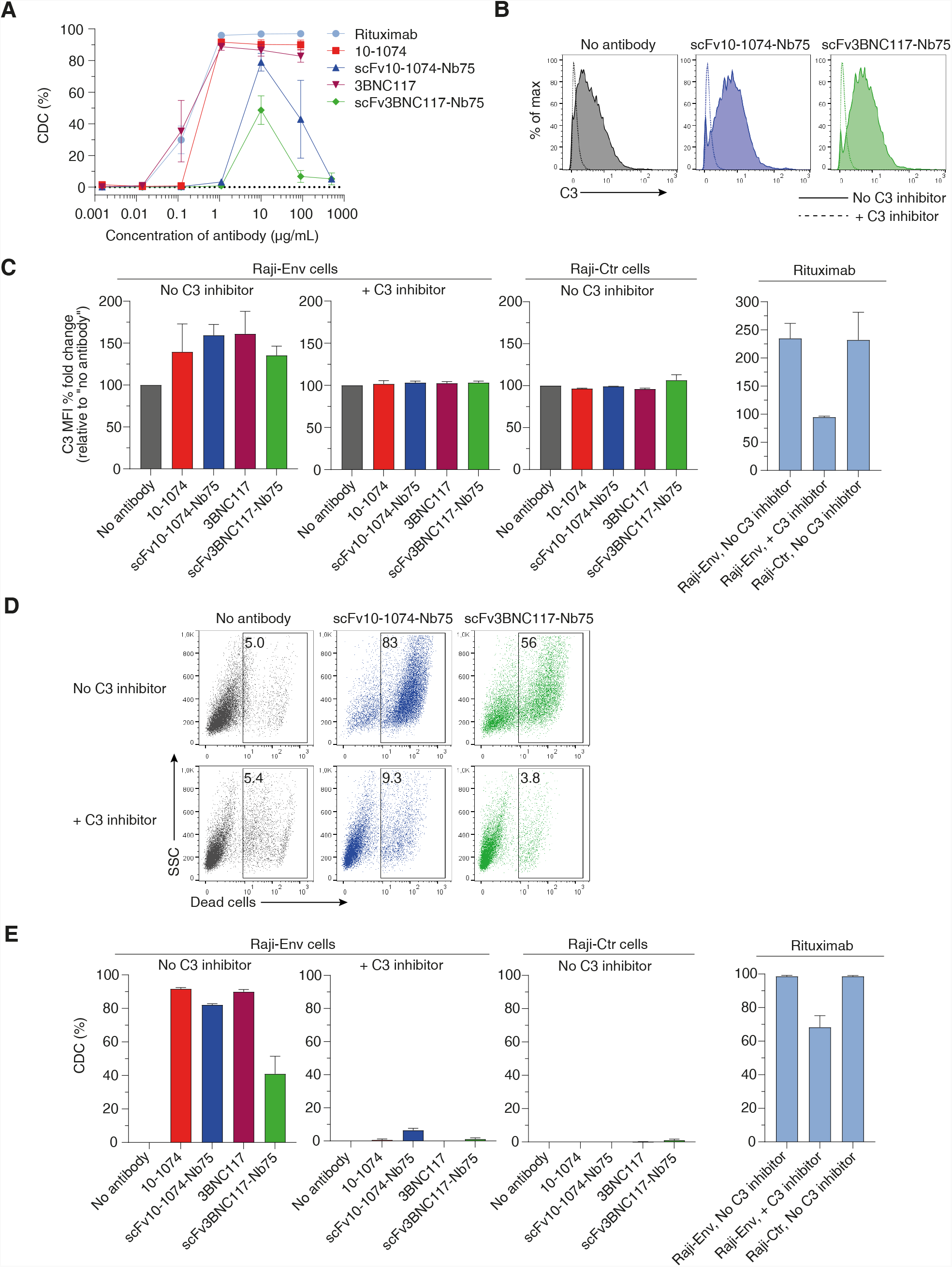
Induction of C3 deposition and CDC of Env-expressing cells by anti-HIV BiCEs. A Raji-Env cells were incubated with NHS and the indicated concentrations of antibodies or BiCEs. 24 h later, cell death was determined by flow cytometry. CDC was calculated as the relative % of dead cells compared to no antibody treatment. n = 3 independent experiments, each with a mix of 3 donor serum. B Raji-Env cells were incubated with NHS and the indicated antibodies or BiCEs and with or without a C3 inhibitor (hC3Nb2). Surface levels of C3 was measured after 1 h by flow cytometry. One representative experiment is shown. C C3 deposition by the indicated antibodies or BiCEs. Results are expressed as the C3 MFI % fold change compared to no antibody treatment. n = 3 donor serum. D Raji-Env cells were incubated with NHS and the indicated antibodies or BiCEs and with or without a C3 inhibitor. Cell death was measured 24 h later by flow cytometry. The numbers indicate the % of dead cells. One representative experiment is shown. D CDC triggered by the indicated antibodies or BiCEs. CDC was calculated as the relative % of dead cells compared to no antibody treatment. n = 3 donor serum. All data are presented as mean ± SEM.

To assess complement activation by anti-HIV BiCEs, we measured the deposition of C3 on cells after 1 h and cell death after 24 h by flow cytometry. Cells cultured in presence of a C3 inhibitor (hC3Nb2), a recently discovered nanobody inhibiting C3 cleavage, were used as negative control (Pedersen *et al*, 2020). C3 deposition was increased on Raji-Env cells in presence of anti-HIV BiCEs compared to no antibody treatment (Figure 3B-C). In presence of C3 inhibitor, the C3 deposition was absent on Raji-Env cells (Figure 3B-C). Furthermore, complement activation was specific for Env expressing cells as C3 deposition was not observed on Raji-Ctr cells (Figure 3C). Additionally, the anti-HIV BiCEs triggered CDC of Raji-Env cells (Figure 3D). ScFv10-1074-Nb75 resulted in a mean of 80% of cell death after 24 h incubation, almost equivalent to CDC caused by 10-1074 and 3BNC117 (Figure 3E). ScFv3BNC117-Nb75 appeared less potent, causing a mean of 40% of cell death (Figure 3E). Moreover, no CDC was observed in presence of C3 inhibitor or with Raji-Ctr cells (Figure 3E). Together these results show that anti-HIV BiCEs are highly capable of recognizing and mediating complement activation and lysis of Env-expressing cells, with scFv10-1074-Nb75 being nearly as potent as full-length bNAbs.

### Complement-mediated anti-HIV effects initiated by anti-HIV BiCEs

Next, we sought to examine whether the anti-HIV BiCEs were able to induce complement activation and subsequent lysis of HIV-1-infected primary CD4 T cells. We used a laboratory-adapted R5-tropic HIV reporter NL4-3 strain (HIV-1_NL4-3-eGFP_), allowing us to measure HIV-infection by eGFP detection. First, we measured C3 deposition mediated by 10-1074 and scFv10-1074-Nb75 by flow cytometry to determine the most suitable concentration to use in subsequent assays with infected primary CD4 T cells. At the highest concentration of antibody or BiCE (30 µg/mL), C3 deposition on cells reached a saturation plateau (Figure S1).

We then assessed complement activation and killing of infected cells triggered by anti-HIV antibodies or BiCEs over time. Primary CD4 T cells were infected with HIV-1_NL4-3-eGFP_ for 24 h. Infected cells were not sorted from uninfected cells, which enabled a specificity evaluation of complement activation on eGFP^-^ cells (uninfected). An HIV integrase inhibitor (raltegravir) was added to the cultures, to prevent viral spread to uninfected cells, precluding that reduction in infection was caused by antibody-mediated neutralization of virus. Cells were cultured in presence of autologous NHS and anti-HIV antibody or BiCE. C3 deposition and CDC were measured by flow cytometry. CDC was quantified as the elimination of eGFP^+^ cells. Cells were analyzed after 1.5 h, 24 h, 3- and 6 days incubation with antibody or BiCE. We observed a significant increase in C3 deposition on infected cells in presence of scFv10-1074-Nb75 and 10-1074 compared to no antibody treatment (Figure 4A-B). Surprisingly, no increase in C3 deposition was detected for scFv3BNC117-Nb75 and 3BNC117 (Figure 4A-B). When adding the two BiCEs or the two full-length antibodies together, an increase in C3 deposition was observed. Combination of antibodies or BiCEs targeting different epitopes likely cause deposition of more antibodies or BiCEs on the cell surface, resulting in increased binding of C1q (Figure 4A-B). As expected, we did not observe any complement activation of eGFP^-^ cells (Figure 4A-B). Moreover, the frequency of eGFP^+^ cells decreased over time in all conditions (Figure 4C + S2). However, in presence of anti-HIV antibody or BiCE, the elimination of eGFP^+^ was accelerated over time (Figure 4C), with the difference between anti-HIV antibodies or BiCEs and no antibody treatment reaching statistical significance at day 6 (Figure 4D). The presence of complement and anti-HIV antibodies or BiCEs did not affect survival of eGFP^-^ cells, indicating that decrement of infected cells was mediated by antibody-dependent complement activation (Figure S2). Overall, these results suggest that our developed anti-HIV BiCEs induce complement-mediated killing, almost at the same level of full-length anti-HIV antibodies.

**Figure 4.**
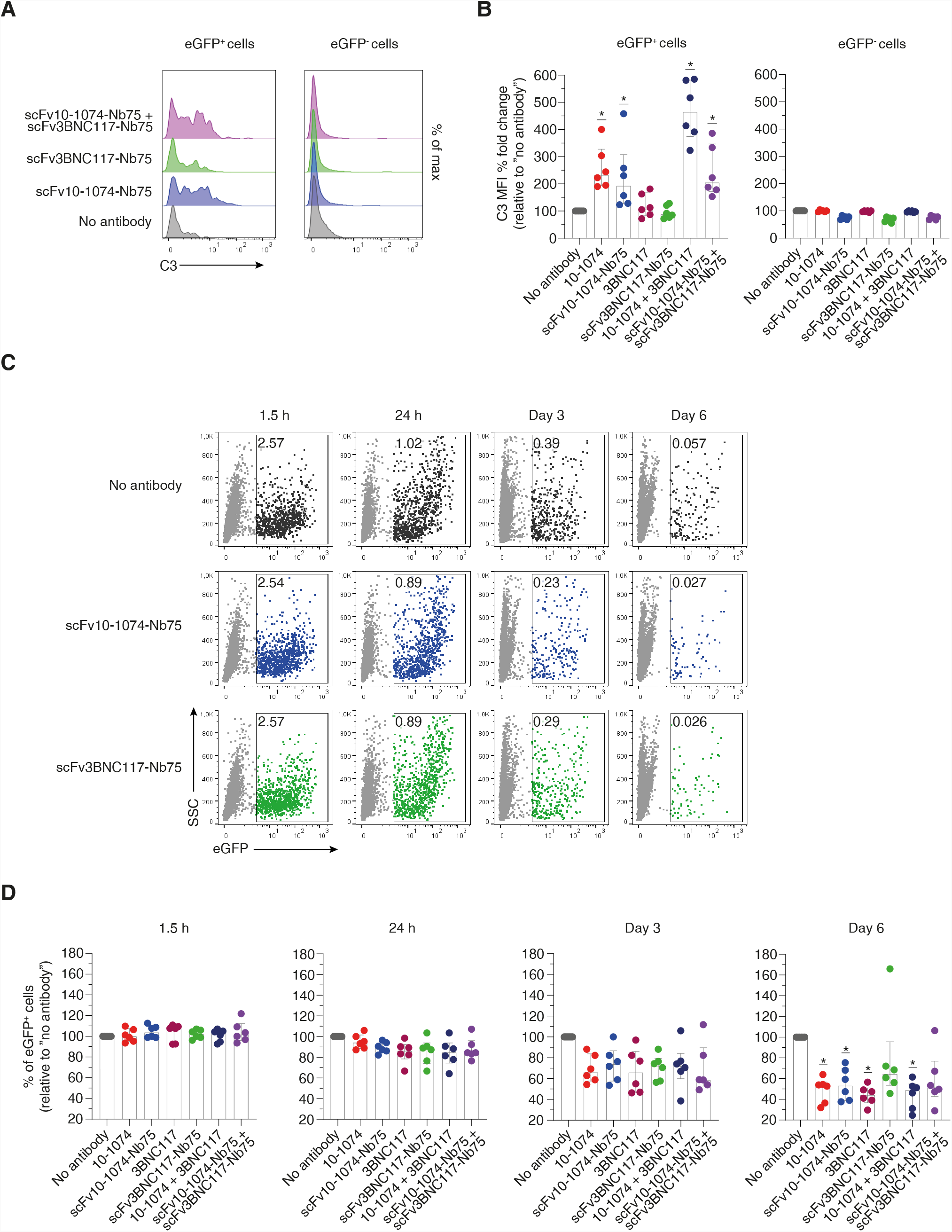
Anti-HIV-1 BiCEs increase complement deposition and elimination of HIV-infected cells. A Primary CD4 T cells infected with HIV-1_NL4-3-eGFP_ and uninfected CD4 T cells were incubated with NHS and the indicated antibodies or BiCEs in presence of an HIV-inhibitor (raltegravir). Surface levels of C3 was measured after 24 h by flow cytometry. One representative donor is shown. B C3 deposition by the indicated antibodies or BiCEs. Results are expressed as the C3 MFI % fold change compared to no antibody treatment. n = 6 donors. C Primary CD4 T cells infected with HIV-1_NL4-3-eGFP_ and uninfected CD4 T cells were incubated with NHS and the indicated antibodies or BiCEs in presence of an HIV-inhibitor. The frequency of eGFP+ cells were analyzed by flow cytometry at the indicated timepoints. The numbers indicate the % of eGFP+ cells. One rep-resentative donor is shown. D Induction of eGFP+ elimination over time by the indicated antibodies or BiCEs. Results are expressed as the relative % of eGFP+ cells compared to no antibody treatment. n = 6 donors. All data are presented as median with interquartile range. Significance was determined by comparing each antibody or BiCE to no antibody treatment. Only significant differences are shown; *P <0.05, Wilcoxon signed-rank test.

## Discussion

While the attempt to approach an HIV cure continues, various strategies are under evaluation. We aimed to assess a potential novel therapeutic modality, based on nanobody-mediated complement recruitment to HIV-infected cells. In this study, we characterized two anti-HIV BiCEs, designated scFv10-1074-Nb75 and scFv3BNC117-Nb75, each composed of a nanobody binding C1q and a bNAb scFv binding the HIV Env. With the development of our anti-HIV BiCEs, we show that C1q can be recruited directly and very specific to the surface of Env-expressing cells. Furthermore, we observe cell death of Env-expressing cells and accelerated elimination of HIV-infected cells while uninfected cells are left unharmed, suggesting CDC as the underlying mechanism of action. Thus, we have accomplished the development of a potential novel therapeutic approach to trigger complement activation and killing of HIV-infected cells.

We establish that anti-HIV BiCEs are highly capable of neutralizing free HIV-1 virions. Furthermore, we observe a minor reduction in neutralizing potency by the two anti-HIV BiCEs compared to bNAbs. The reduction in potency of the BiCEs is consistent with other studies determining the neutralizing potency of different bNAb scFvs and the matched IgG, in which the difference in IC_50_ values range in average between 4-to 13-fold (van Dorsten *et al*, 2020). This reduction in potency presumably mirrors the presence of two antigen-binding sites in full-length IgG compared to one binding site in the scFv part of the BiCEs.

Using Env-expressing Raji cells, resembling HIV-infected cells on their surface, we examine the specificity of our two anti-HIV BiCEs as well as complement-mediated effects. We observe that the levels of complement activation triggered by our anti-HIV BiCEs resemble the capacities of the respective full-length bNAb. Moreover, complement activation is accompanied by increased CDC, with scFv10-1074-Nb75 showing similar levels of CDC as full-length 10-1074. In contrast to most conventional antibodies, the dose-dependent effects of anti-HIV BiCEs decrease at high concentrations. This is in line with other bispecific antibodies used in oncology, in which different variables such as the expression of target proteins, the ratio between bispecific antibodies and its target proteins, and the binding affinity of the bispecific antibody for its specific target proteins impact the efficacy window of the bispecific antibody (Betts & van der Graaf, 2020). Moreover, anti-HIV BICEs may have a shorter half-life due to the lack of an Fc domain, precluding recycling by the neonatal Fc receptor (Roopenian & Akilesh, 2007). However, the pharmacokinetics of anti-HIV BiCEs remains to be examined.

In the current study, we also examine the killing of HIV-infected primary cells triggered by anti-HIV BiCEs. The complement-mediated effects of scFv10-1074-Nb75 and 10-1074 on HIV-infected primary CD4 T cells are similar to those on Raji-Env cells. In contrast to earlier findings, however, only limited complement deposition by scFv3BNC117-Nb75 and 3BNC117 is detected (Dufloo *et al*., 2020). There may be notable distinctions in the HIV Env structure and density of the YU-2 Env expressed by Raji-Env cells versus the NL4-3 Env expressed by the HIV-infected primary CD4 T cells that likely affect the sensitivity to antibodies and potentially explain the differences observed. Despite low-level of C3 deposition, scFv3BNC117-Nb75 and 3BNC117 accelerate the elimination of HIV-infected primary cells similar to that of scFv10-1074-Nb75 and 10-1074.

We demonstrate that anti-HIV BiCEs and antibodies induce CDC of Raji cells within 24 hours, consistent with a previous study of anti-HIV antibodies (Dufloo *et al*., 2020). However, we observe slower complement-mediated effects of HIV-infected primary CD4 T cells. Primary HIV-infected cells are different from the B cell lymphoma Raji-Env cell line in a number of respects. Slower elimination of infected cells is likely explained by the presence of CD59 on primary CD4 T cells, a key regulatory complement protein, preventing MAC formations on cell surfaces (Bajic *et al*., 2015). Moreover, CD59 is incorporated into the viral lipid envelope during viral budding from host cells, providing resistance to CDC of viral particles (Saifuddin *et al*, 1995). In contrast, Raji cells are highly susceptible to CDC as they have low-level expression of CD59 (Lara *et al*, 2021; Tone *et al*, 1999; Wang *et al*, 2010). Several reports show that knock-out of CD59 restores the sensitivity of cells and HIV virions to complement-mediated effects (Dufloo *et al*., 2020; Lan *et al*, 2014; Yang *et al*, 2015). In addition, viral proteins, including Vpu and Nef, protect infected cells from complement- and antibody-mediated lysis by down-regulating and modulating envelope surface levels and conformations, resulting in limited antibody recognition (Alvarez *et al*, 2014; Arias *et al*, 2014; Dufloo *et al*., 2020; Pham *et al*, 2014).

A key element in adaptive immune responses against HIV-1 is antibody-mediated clearance of HIV and HIV-infected cells. More specifically, Fc-domains of bNAbs engage with Fc receptors (FcRs) present on immune cells, facilitating antibody-dependent cellular cytotoxicity (ADCC) and antibody-dependent cellular phagocytosis (ADCP) against HIV and HIV-infected cells (Bruel *et al*, 2016; Musich *et al*, 2017; von Bredow *et al*, 2016). Anti-HIV BiCEs lack an Fc-domain, precluding critical adaptive immune responses like ADCC and ADCP. However, in vitro studies show that complement-dependent adaptive immune responses, including complement-dependent cellular cytotoxicity (CDCC) and complement-dependent cellular phagocytosis (CDCP) mediate efficient clearance of target cells similar to those of the FcR-dependent effector functions (Lee *et al*, 2017). Anti-HIV BiCEs may be capable of inducing CDCC and CDCP, however, this remains to be explored.

We compared the anti-HIV BiCEs with their respective full-length bNAb in the experimental settings. In vitro and ex vivo complement activation assays show that anti-HIV BiCEs are either equivalent potent or slightly less potent compared to the respective parental bNAb. However, the approximately 3-fold smaller size of BiCEs compared to full-length bNAbs may enable increased tissue penetrance into the densely packed lymphoid tissues harboring HIV-infection such as lymph nodes, spleen, and gut (Chen & Dimitrov, 2009). Moreover, the smaller size of BiCEs may allow competition for epitope sites in close proximity, such as the V3 loop and CD4 binding site of HIV Env (Labrijn *et al*, 2003).

The options of developing BiCEs with different targeting arms are numerous. This enables combination therapy with binding of multiple epitopes of the HIV Env. Previous publications of clinical trials showed that combination therapy with the bNAbs 3BNC117 and 10-1074 resulted in viral suppression and prevented viral escape mutations over a median period of 21 weeks after analytical treatment interruption (Mendoza *et al*, 2018). Here, we report that ex vivo combination of scFv10-1074-Nb75 and scFv3BNC117-Nb75 induce elimination of HIV-infected primary cells almost in line with combination of 3BNC117 and 10-1074.

The major barrier to cure HIV is the persistence of a latent reservoir despite successful cART treatment (Pitman *et al*, 2018). Studies have showed that activation of latent provirus alone is not enough to eradicate the virus, suggesting that treatment with latency reversal agents should be followed by boosting of the immune response to destroy the infected cells (Kim *et al*, 2018). With killing mediated by our two anti-HIV BiCEs, treatment with anti-HIV BiCEs could be implemented as a part of the “kick and kill” strategy in combination with latency reversal agents. Since the nanobody-binding domain of BiCEs, recruit C1q and induce complement activation, anti-HIV BiCEs could serve as “the kill” in the “kick and kill” strategy without relying on potential exhausted immune effector cells.

Additionally, treatment with anti-HIV BiCEs may benefit immunocompromised patients, as anti-HIV BiCEs are able to elicit their effector functions without requiring immune effector cells. Moreover, classical complement pathway components are stably expressed in circulation throughout life whereas immune cells are impaired with age, highlighting the advantageous use of BiCEs in individuals with diminished immune capacity (Gaya da Costa *et al*, 2018; Haynes & Maue, 2009; Wenisch *et al*, 2000).

In conclusion, we have developed a bispecific complement engager with great propensity for complement engagement. Our results demonstrate proof-of-concept that these anti-HIV BiCEs may offer new and highly specific opportunity to utilize complement-mediated killing of HIV-infected cells in an HIV cure setting.

## Materials and methods

### Cells

HEK293T cells and TZM-bl cells (NIH HIV Reagent Program #ARP-8129) were cultured in complete DMEM (cDMEM) that is: DMEM (Biowest) + 10% heat-inactivated fetal bovine serum (HI-FBS) (Biowest) + 100 U/mL Penicillin G and Streptomycin (P/S) (Biowest). All cells were incubated at 37°C and 5% CO_2_ in a humidified environment.

Raji cells were cultured in complete RPMI (cRPMI) that is: RPMI 1640 (Biowest) + 10% HI-FBS + 100 U/mL P/S. Engineered Raji Env-expressing cells have been previously described and were kindly provided by J. Dufloo and O. Schwartz from Institute Pasteur (Dufloo *et al*., 2020). Briefly, Raji cells were spinoculated with a retroviral vector carrying YU-2 Env (pMX-YU2 ENVΔCT-GFP-Puro^R^). Transduced cells (GFP^+^) were cultured in presence of 1 µg/mL puromycin (Gibco). In our lab, co-expression of GFP and Env was validated and determined to be 94% (data shown in Figure S3).

Peripheral blood mononuclear cells (PBMCs) and serum were obtained from peripheral blood of healthy human donors from the Department of Clinical Immunology, Blood Bank, at Aarhus University Hospital, according to local ethical guidelines. PBMCs were isolated using Ficoll-Paque PLUS density gradient separation (GE Healthcare).

CD4 T cells were enriched from PBMCs by negative immunomagnetic selection using CD4+ T cell isolation kit (Miltenyi Biotec), according to the manufacturer’s protocol. The CD4 T cells were cultured in cRPMI. Cells were activated for 3 days in presence of 100 U/mL IL-2 (Gibco) and 1 µg/mL PHA (Remel).

### Viruses and infection

Laboratory adapted HIV-1_YU-2_ clone was obtained from the NIH HIV Reagent Program (Cat. no. #ARP-1350). Full-length HIV-1_NL4-3-eGFP_ reporter virus was produced using a pBR-HIV-1 M NL4-3 92TH14-12 plasmid. The plasmid was generated and has been previously described by A. Papkalla and F. Kirchhoff (Papkalla *et al*, 2002). The production of virus is outlined below. The plasmids were transfected into HEK293T cells using DNA:PEI ratio of 1:4 for expression and virus production. After 72 hours, the supernatant containing the virus was harvested and filtered, and stored at −80°C.

Prior to infection of activated CD4 T cells, the PHA-containing medium was replaced with fresh cRPMI supplemented with 100 U/mL IL-2. HIV-1_NL4-3-eGFP_ was added to the cells at a multiplicity of infection (MOI) of 0.1. Cells were spinoculated at 1250 x g with virus for 2 h and cultured overnight. Next day, media was replaced to remove unbound virus before performing further experiments.

### Antibodies and recombinant proteins

Full-length anti-Env monoclonal antibodies 10-1074 and 3BNC117 were used (Mouquet *et al*., 2012; Scheid *et al*., 2011). Additionally, the anti-CD20 antibody Rituximab (Rixathon; Sandoz) was used.

scFv10-1074 was produced by basic cloning techniques and expression of protein. Briefly, the scFv10-1074 DNA inserts were cloned into a pcDNA 3.1(+). The cloned pcDNA3.1(+)-scFv10-1074 was transfected into HEK293T cells for expression and protein production. The supernatant containing the scFv-10-1074 was harvested and filtered, and stored at −80°C. The generated scFv10-1074 protein contained a biotin tag for purification.

ScFv10-1074-Nb75 was constructed as V_L_-(GGGGS)_3_-V_H_-(GGGGS)_3_-C1qNb75-6xHIStag and scFv3BNC117-Nb75 was constructed as V_H_-(GGGGS)_3_-V_L_-(GGGGS)_3_-C1qNb75-6xHIStag. For these constructs, the V_H_ and V_L_ sequence was that of full-length 10-1074 and 3BNC117 antibody, and C1qNb75 has been previously described by NS Laursen and GR Andersen (Laursen *et al*., 2020; Mouquet *et al*., 2012; Scheid *et al*., 2011).

The BiCE constructs were cloned into pcDNA3.1 and expressed in HEK293F cells using DNA:PEI ratio of 1:2. Conditioned media was harvested after 5 days of incubation and filtered. Tris pH 8.0 was added to 50 mM final concentration. Supernatant was loaded on HisTrap FF and washed to baseline with 0.1 M Tris pH 8.0, 500 mM NaCl, 30 mM imidazole. Protein was eluted with wash buffer supplemented with 400 mM imidazole. Finally, eluate was concentrated and loaded on superdex75 inc. (24ml) equilibrated in 20 mM HEPES, 150 NaCl pH7.3. Complex formation was tested by SEC using a superdex75 inc. (24ml) equilibrated in 20 mM HEPES, 150 NaCl pH7.3. BiCEs and C1qGH were mixed using 1.2-fold molar excess of C1qGH.

### Antibody neutralization assay

Neutralization by anti-HIV antibodies and BiCEs was assessed through infection of HIV reporter TZM-bl cells with a laboratory-adapted HIV-1_YU-2_ clone. Three-fold serial dilutions was made of each antibody and BiCE in cDMEM. All samples were added in triplicate wells to a 96-well culture plate. Virus was added at MOI = 0.1 to each well and incubated for 1 h at 37°C. Then, TZM-bl cells were added to each well and incubated for 48 h at 37°C in presence with 12.5 μg/mL DEAE-Dextran (Sigma-Aldrich). Prior to data acquisition, cells were lysed with 0.5% NP40 (BDH Laboratory supplies) for 45 min. at room temperature (RT). Britelite plus (Perkin Elmer) was added to each well. The solution was transferred to a white 96-well Optiplate (PerkinElmer), and the relative luminescence unit (RLU) was measured with a FLUOstar Omega plate reader (BMG Labtech). RLU for background control wells without virus was subtracted from all other wells. Infection in virus control wells was set to 100%.

### Expression of HIV-Env on Raji cell surfaces

Raji-Env and Raji-Ctr cells (1×10^5^) was washed in PBS containing 2% HI-FBS, and pre-incubated with Human TruStain FcX blocker (BioLegend) for 5 min at RT. Subsequently, the cells were incubated with or without 30 µg/mL of anti-HIV antibodies for 45 min at 4 °C. Cells were prepared for flow cytometry by washing once with PBS containing 2% HI-FBS followed by incubation with Brilliant Violet 421 anti-human IgG Fc Antibody (clone M1310G05) (Biolegend) or Brilliant Violet 421 Streptavidin (BioLegend) (gating strategy shown in Figure S4).

### Complement activation assay: Raji cells

Raji-Env and Raji-Ctr cells (5×10^4^) were resuspended in 100 µL RPMI containing 25% NHS from healthy donors and 100 U/mL P/S. Cells were seeded in a 96-well culture plate and incubated for 1 h and 24 h at 37°C with or without 10 µg/mL antibody or BiCE (unless otherwise stated), and with or without a C3 nanobody inhibitor (hC3Nb2) using C3:hC3Nb2 molar ratio of 1:4. HC3Nb2 has been previously described by H. Pedersen and GR Andersen (Pedersen *et al*., 2020). After 1 h and 24 h, the cells were prepared for flow cytometry. To measure CDC, cells were stained with a live/dead marker: Zombie Violet Fixable Viability Stain (Biolegend) for 30 min at 4°C. CDC was determined using the following formula: 100 x (% of dead cells with antibody - % of dead cells without antibody) / (100 - % of dead cells without antibody). Negative values were set to zero. Second, to measure complement deposition, cells were pre-incubated with Human TruStain FcX blocker (BioLegend) for 5 min at RT, followed by incubation with APC anti-complement C3b/iC3b Antibody (clone 3E7/C3b) (BioLegend) or APC Mouse IgG1, κ Isotype Ctrl Antibody (clone MOPC-21) (BioLegend) for 20 min at 4°C (gating strategy shown in Figure S4).

### Complement activation assay: CD4 T cells

CD4 T cells (1×10^5^) were resuspended in 100 µL RPMI containing 50% autologous NHS, 100 U/mL P/S and 100 U/mL IL-2. Moreover, the cells were maintained in presence of 500 nM Raltegravir (Merck). Cells were seeded in a 96-well culture plate and incubated with or without 30 µg/mL antibody or BiCE (unless otherwise stated) for 1 h, 24 h, 3- and 6 days at 37°C. To measure complement deposition and CDC, the cells were prepared for flow cytometry. Complement deposition was measured as described above. Cells were fixed with 1% PFA (Thermo Scientific) for 15 min at 4°C before complement deposition and eGFP expression were analyzed (gating strategy shown in Figure S4). Elimination of eGFP^+^ cells was used as a measure of CDC, calculated with the following formula: 100 x (% of eGFP^+^ cells without antibody - % of eGFP^+^ cells with antibody) / (% of eGFP^+^ cells without antibody). Negative values were set to zero.

### Data processing and statistical analysis

Miltenyi Biotec MACSquant16 flow cytometer was used to acquire data by flow cytometry. Data was analyzed using FlowJo (version 10.8.1). All data processing and transformations was carried out in Microsoft Excel 365. GraphPad Prism (version 9.3.1) was used for graphing and statistical analysis. When relevant, statistical analysis was performed between groups using non-parametric Wilcoxon signed rank test. Only significant differences (p ≤ 0.05) are highlighted in each graph.

## Data availability

This study includes no data deposited in external repositories.

## Acknowledgement

We thank members of the Infectious Diseases Research Unit for scientific discussion and help. We thank Emma Falling Iversen and Kathrine Kjær for virus production, and Christina Valbirk Konrad for the generation of scFv10-1074.

We thank Dr Oliver Schwarts (Institute Pasteur) for kindly providing the Raji-Env and Raji-Ctr cells. The following reagent was obtained through the NIH HIV Reagent Program, Division of AIDS, NIAID, NIH: TZM-bl Cells, ARP-8129, contributed by Dr. John C. Kappes and Dr. Xiaoyun Wu; and Human Immunodeficiency Virus 1 (HIV-1) YU2 Infectious Molecular Clone, ARP-1350, contributed by Dr. Beatrice Hahn and Dr. George M. Shaw. Figure 1A was created with BioRender.com.

MT was supported by grant from the Novo Nordisk Foundation (grant number NNF19OC0054577) and the Danish Independent Research Fund (grant number 9039-00039B). AHFA was supported by a scholarship from the Danish Independent Research Fund (grant number 1029-00004B).

## Author contributions

Conceptualization: MLP, DVP, NSL, AHFA, MT

Funding acquisition: AHFA, MT

Investigation: MLP, DVP

Methodology: MLP, DVP, MBLW, HGO, NSL, AHFA, MT

Resources: OSS, LØ, AHFA, MT

Supervision: AHFA, MT Visualization: MLP, DVP

Writing – original draft: MLP, DVP, AHFA, MT

Writing – review & editing: MLP, DVP, MBLW, HGO, OSS, LØ, NSL, AHFA, MT

## Supplementary figure legends

**Figure S1.**
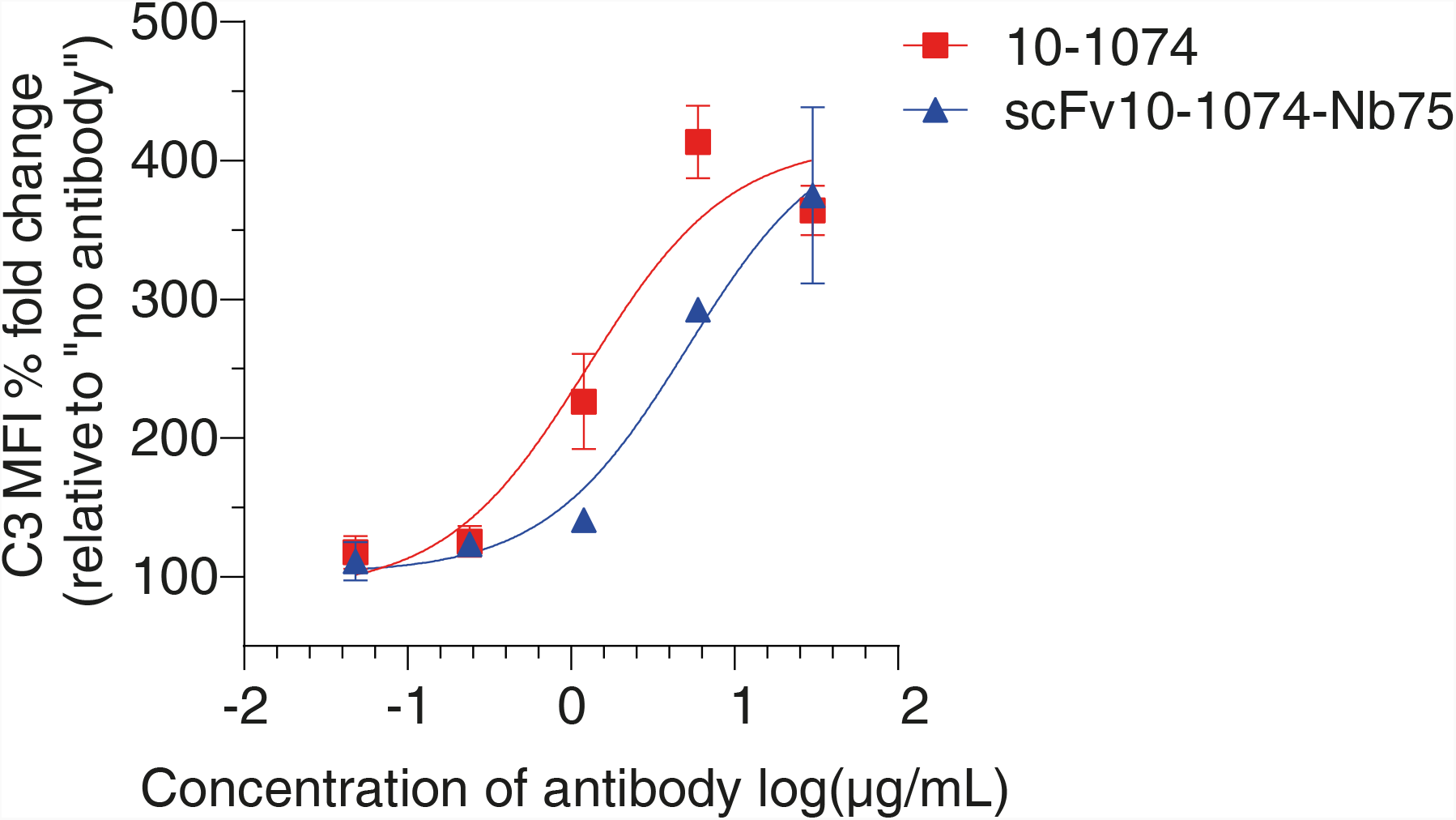
Concentration-dependent C3 deposition of infected primary CD4 T cells. Primary CD4 T cells infected with HIV-1_NL4-3-eGFP_ were incubated with NHS and the indicated concentrations of antibodies or BiCEs. After 24 h, surface levels of C3 were measured by flow cytometry. n = 2 donors. Data is presented as median with range.

**Figure S2.**
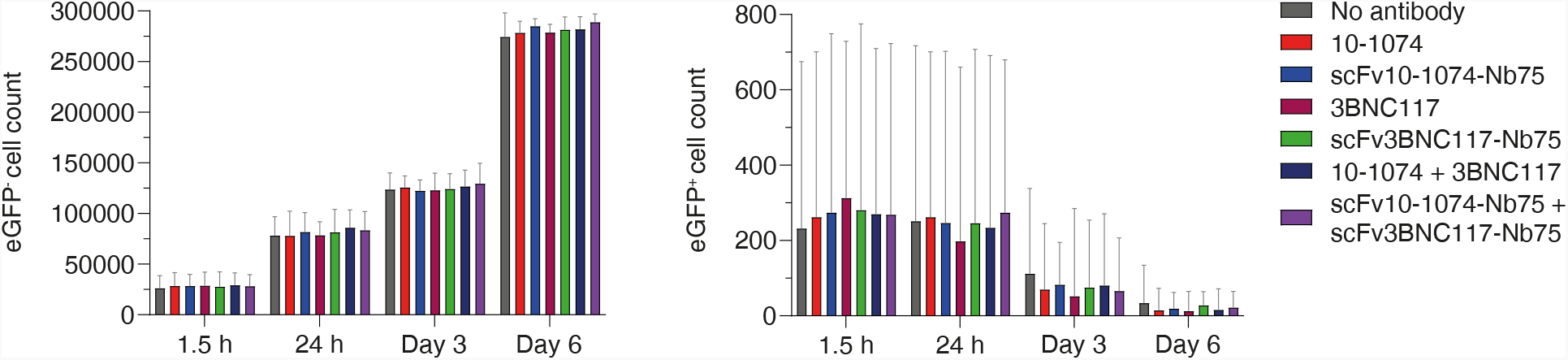
Presence of NHS and anti-HIV antibodies or BiCEs do not affect uninfected cells. Primary CD4 T cells infected with HIV-1_NL4-3-eGFP_ and uninfected CD4 T cells were incubated with NHS and the indicated antibodies or BiCEs in presence of an HIV-inhibitor. The cell count was assessed by flow cytometry at the indicated timepoints. n = 4 donors. Data is presented as median with interquartile range.

**Figure S3.**
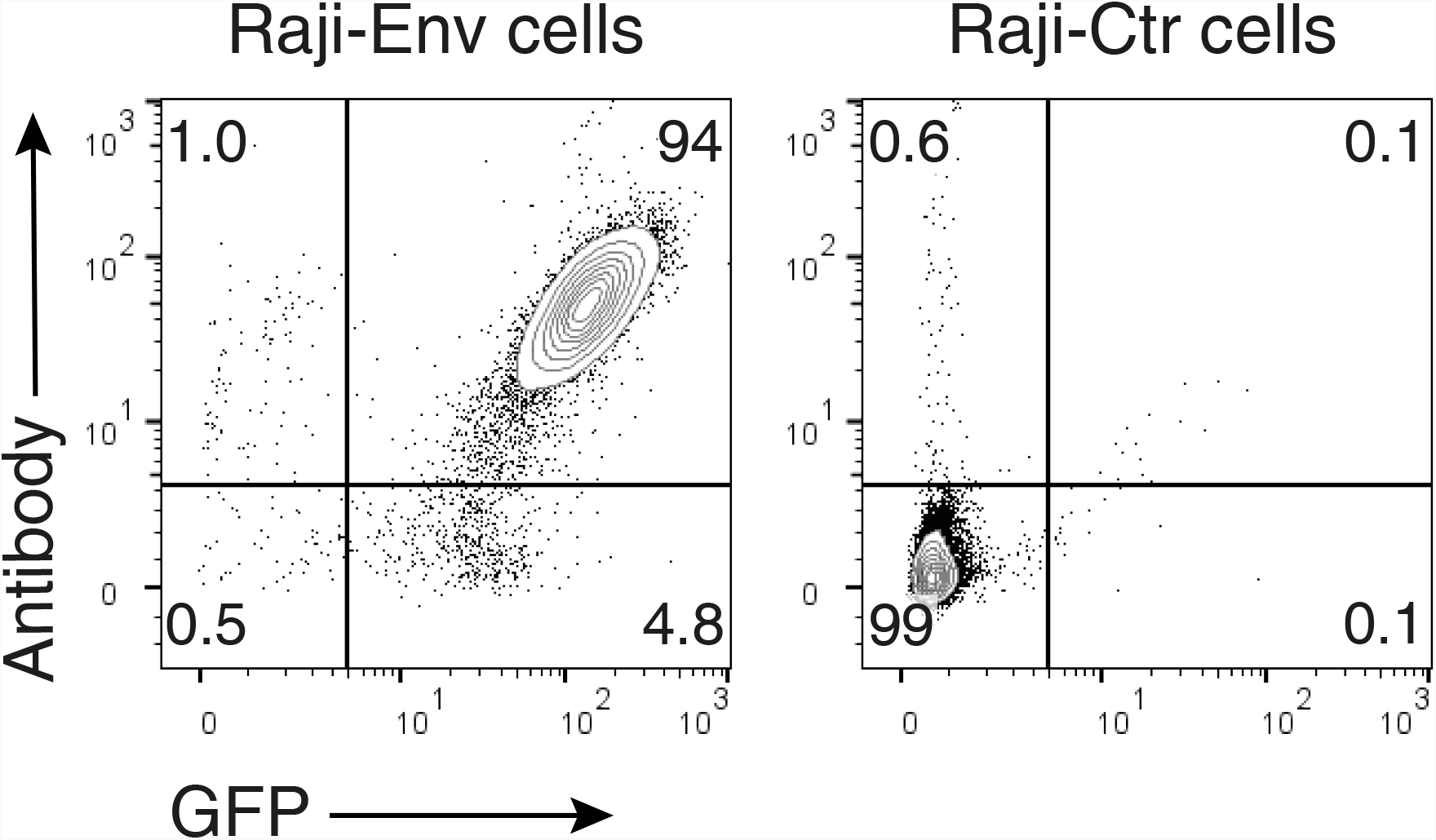
Co-expression of GFP and Env on Raji cells. Raji cells expressing HIV-1_YU-2_ envelope (Raji-Env) or Raji control cells (Raji-Ctr) were incubated with anti-Env monoclonal antibody 10-1074. After 45 min, binding of 10-1074 to HIV-Env on cells were assessed by flow cytometry using a BV421-conjugated anti-IgG Fc antibody. The numbers indicate the percentage of cells.

**Figure S4.**
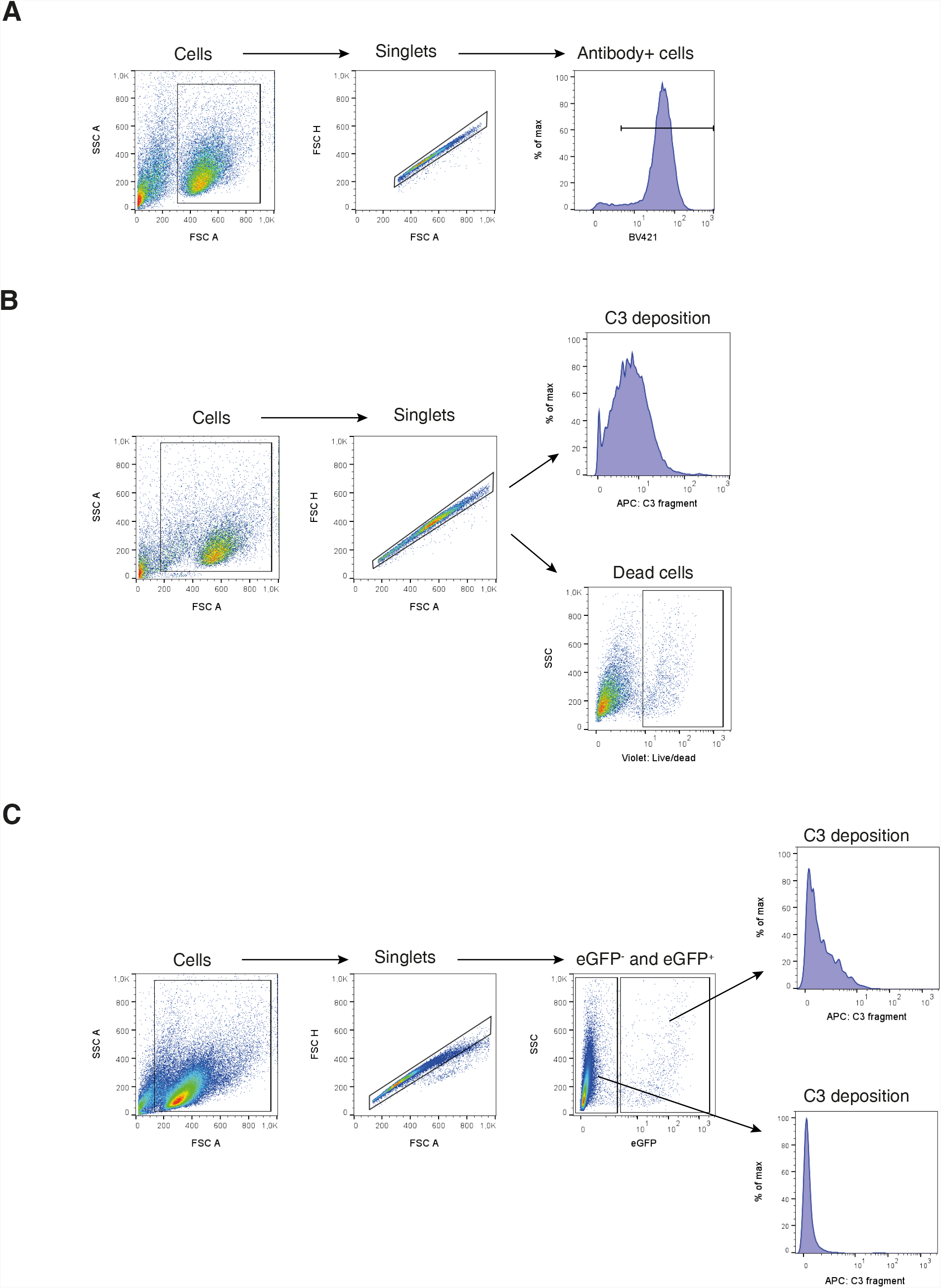
Gating strategy for flow cytometry assays. A Gating strategy for anti-HIV antibody surface stain of Raji cells. BV421-conjugated anti-IgG Fc or BV421-conjugated streptavidin were used to discriminate cells with bound anti-HIV antibody. B Gating strategy for complement activation assays of Raji cells. APC-conjugated anti-C3 antibody and zombie violet live/dead marker were used to differentiate cells with deposited C3 and dead cells, respectively. C Gating strategy for complement activation assays of primary CD4 T cells. EGFP expression was used to distinguish HIV-infected cells. APC-conjugated anti-C3 antibody was used to discriminate cells with deposited C3.

